# Temporal gating of synaptic competition in the lateral amygdala by cannabinoid receptor modulation of the thalamic input

**DOI:** 10.1101/526624

**Authors:** Ana Drumond, Natália Madeira, Rosalina Fonseca

## Abstract

The acquisition of fear memories involves plasticity of the thalamic and cortical pathways to the lateral amygdala (LA). The maintenance of synaptic plasticity requires the interplay between input-specific synaptic tags and the allocation of plasticity-related proteins (PRPs). Based on this interplay, weakly activated synapses can express long-lasting synaptic plasticity by cooperation with strongly activated ones. Increasing the number of activated synapses can shift cooperation to competition. Synaptic cooperation and competition can determine whether two events, separated in time, are linked or selected. The rules that determine whether synapses cooperate or compete are unknown. We found that synaptic cooperation and competition, in the LA, are determined by the temporal sequence of cortical and thalamic stimulation and that the strength of the synaptic tag is modulated by the endocannabinoid signalling. This modulation is particularly effective in thalamic synapses, suggesting a critical role of endocannabinoids in restricting thalamic plasticity. Also, we found that PRPs availability is modulated by the action-potential firing of neurons, shifting competition to cooperation. Our data present the first evidence that pre-synaptic modulation of synaptic activation, by the cannabinoid signalling, function as a temporal gating mechanism limiting synaptic cooperation and competition.

## INTRODUCTION

Learning is a key process allowing individuals to adapt to environmental challenges (McKenzie and Eichenbaum, 2011). Cellular models of memory, such as long-term potentiation (LTP), share common principles with memory consolidation (Bliss and Collingridge, 1993). As for memory, not all events are maintained in long-lasting forms of LTP. Depending on the strength of synaptic activation, LTP induction can set a local, input-specific, synaptic tag or it can also induce the synthesis of PRPs (Frey and Morris, 1997; Sajikumar and Frey, 2004). If protein synthesis is triggered, LTP is maintained. Since the setting of the “synaptic tag” and the capture of PRPs are two independent processes, then induction of a maintained form of LTP in one set of synapses can stabilize a transient form of LTP induced in a second independent set of synapses (Redondo and Morris, 2011). This cooperative maintenance allows neurons to integrate events that occur within large time windows, ranging from 30 to 60 minutes (Frey and Morris, 1998; Sajikumar et al., 2007). If PRPs are limited, for example by blocking protein-synthesis, or by increasing the pool of activated synapses, then synaptic competition is observed (Fonseca et al., 2004; Govindarajan et al., 2011; Sajikumar et al., 2014). In this working model, synaptic cooperation and competition are determined by a balance between PRPs availability (source) and the strength of the tag (sink). In turn, the tag strength is a combination of the number of activated synapses and the ability that each synapse has in capturing PRPs (Fonseca, 2012; Szabó et al., 2016). Previous studies looking at competition in CA1 hippocampal synapses have shown that competition was correlated with the strength of the tag (Fonseca et al., 2004). According to this, synapses were stronger or weaker depending on their relative activation and therefore, winning synapses block the maintenance of LTP in loser synapses. However, this was not observed in a second study, where competition displayed a “winner-take-all” form. In this case, strong and weak activated synapses were similar in their ability to capture PRPs and neither were maintained in a competitive setting (Sajikumar et al., 2014).

Cooperation in the lateral-amygdala follows similar principles as shown for hippocampal synapses. Lateral amygdala pyramidal cells receive input from the auditory cortex (C) and the auditory thalamus (T), a neuronal circuitry that is involved in the acquisition of fear memories (Amano et al., 2010). Lesion studies have shown that the activation of either thalamic or cortical inputs is sufficient to acquire a fear memory, whereas both inputs are necessary to discriminate between fearful and neutral events (Antunes and Moita, 2010). We have previously shown that cortical and thalamic synapses can cooperate by a synaptic tagging and capture mechanism (Fonseca, 2013). Interestingly, thalamic and cortical synapses were not identical in their ability to capture PRPs and maintain LTP by cooperation. Thalamic synapses had a much shorter time window of cooperation and this time-window was restricted by the activation of presynaptic cannabinoid receptors (CB1R). This finding gains further relevance if one considers the established link between endocannabinoid signalling and the acquisition of fear memories (Drumond et al., 2017; Mechoulam and Parker, 2013). Endocannabinoids are synthesised on-demand, triggered by neuronal activation and once synthesised, cross the neuronal membrane and bind to their receptors (type 1 cannabinoid receptor – CB1R – and type 2 cannabinoid receptor – CB2R), resulting in a suppression of neurotransmitter release at excitatory and inhibitory synapses (Kano et al., 2009; Turu and Hunyady, 2010). Their degradation by the Fatty acid amide hydrolase (FAAH) and the DAGL (DAG-hydrolizing enzyme) terminate their signalling (Drumond et al., 2017). CB1Rs are highly expressed in the amygdala with regional differences between the lateral, basal and central amygdala (Marsicano and Lutz, 1999). There is compelling evidence that CB1R activation reduces amygdala excitability (Azad et al., 2003; Lutz et al., 2015) and together with a recent report that endocannabinoid synthesis is sensitized by previous negative events (Sumislawski et al., 2011), support the hypothesis that endocannabinoid signalling restricts fear-memory acquisition.

Here, we assessed the temporal rules that determine whether cortical and thalamic synapses interact by cooperation or competition. Further, we tested the role of the endocannabinoid signalling in competition and their contribution in the modulation of cortical and thalamic synaptic plasticity. Our data support the hypothesis that synaptic competition is a cellular mechanism to select the events that are maintained and its temporal restriction may promote memory selectivity.

## MATERIALS AND METHODS

A total of 260 slices prepared from 104 male Sprague-Dawley rats (3-5 week old) were used for electrophysiological recordings. All procedures were approved by the Portuguese Veterinary Office (Direcção Geral de Veterinária e Alimentação – DGAV). Coronal brain slices (350 μm) containing the lateral amygdala were prepared as described previously (Fonseca, 2013). Whole-cell current-clamp synaptic responses were recorded using glass electrodes (7-10MΩ; Harvard apparatus, UK), filled with internal solution containing (in mM): K-gluconate 120, KCl 10, Hepes 15, Mg-ATP 3, Tris-GTP 0.3 Na-phosphocreatine 15, Creatine-Kinase 20U/ml (adjusted to 7.25 pH with KOH, 290mOsm). Putative pyramidal cells were selected by assessing their firing properties in response to steps of current (Supplementary Figure 1A). Voltage-clamp synaptic currents were recorded using 2-3MΩ glass electrodes filled with internal solution. Only cells that had a resting potential of less than −60mV without holding current were taken further into the recordings. Neurons were kept at −70mV to −75mV with a holding current below −0.25nA. In current clamp recordings, the series resistance was monitored throughout the experiment and ranged from 30MΩ-40MΩ; in voltage clamp recordings, series resistance ranged from 10-20MΩ; changes exceeding 25% of the series resistance determined the end of the recording. Electrophysiological data were collected using an RK-400 amplifier (Bio-Logic, France) filtered at 1 kHz and digitized at 10kHz using a Lab-PCI-6014 data acquisition board (National Instruments, Austin, TX) and stored on a PC. Offline data analysis was performed using a customized LabView-program (National Instruments, Austin, TX). To evoke synaptic EPSP, tungsten stimulating electrodes (Science Products, GmbH, Germany) were placed on afferent fibers from the internal capsule (thalamic input) and from the external capsule (cortical input–Supplementary Figure 1A’). Pathway independence was checked by applying two pulses with a 30ms interval to either thalamic or cortical inputs and confirming the absence of crossed pair-pulse facilitation. EPSPs were recorded with a test pulse frequency for each individual pathway of 0.033 Hz. After 20 min of baseline, long-lasting LTP (L-LTP) was induced with a strong tetanic stimulation (25 pulses at a frequency of 100 Hz, repeated five times, with an interval of 3 sec), whereas transient LTP (E-LTP) was induced with a weak tetanic stimulation (25 pulses at a frequency of 100 Hz, repeated three times with an interval of 3 sec). DSE was induced by depolarizing the cell (in voltage-clamp mode) to 0mV during 10 sec.

Drugs were dissolved in DMSO (0.01%) and diluted to achieve the final concentration: Rapamycin (Tocris) 1 μM, AM281 (Sigma) 0.5μM or 1 μM, URB597 (Tocris) 1μM, Picrotoxin (Sigma) 25μM, Win55,212-12 (Tocris) 5μM or 10 μM and SCH-50911 (Tocris) 10 μM. In control experiments, only DMSO was added to the ACSF.

As a measure of synaptic strength, the initial slope of the evoked EPSPs was calculated and expressed as percent changes from the baseline mean. Error bars denote SEM values. For the statistical analysis, LTP values were averaged over 5 min data bins immediately after LTP induction (T Initial – see timeline for each particular pathway and experimental condition) and at the end of the recording (T Final 100-105 minutes). LTP decay was calculated by [(T Initial – T Final)/T Final*100]. Normality was assessed using the Kolmogorov-Smirnov and the Shapiro-Wilk test. Since some of our groups were not normal, group differences were assessed using a non-parametric test (Kruskal–Wallis Test, SPSS software). For the correlation analysis a multiple regression analysis was used (SPSS Software). In Depolarization-suppression of excitation (DSE) experiments, the maximal amplitude of currents evoked by thalamic or cortical stimulation (EPSC) was compared before and after depolarization (Kruskal–Wallis test). For the analysis of the decrease in EPSP slope and changes in PPF, induced by CB1R agonist, we used the Friedman ANOVA (SPSS software) and compared changes before and after agonist application.

## RESULTS

### Stimulation of a second thalamic input leads to competition and destabilization of a previous induced LTP at thalamic synapses

Both thalamic and cortical input afferents to the lateral amygdala can express transient and maintained forms of LTP, depending on the stimulation strength (Fonseca, 2013). Weak tetanic stimulation of thalamic or cortical inputs led to an increase in EPSP slope that decayed to baseline values at the end of the recording (Supplementary Figure 1B/C). If cortical synapses were stimulated with a strong tetanic stimulation, prior to weak thalamic stimulation, thalamic LTP was maintained despite the weak synaptic stimulation (Supplementary Figure 1D). This heterosynaptic cortical-to-thalamic cooperation is dependent on the synthesis of plasticity-related proteins. Bath application of rapamycin, an mTOR pathway inhibitor (Sosanya et al., 2015), led to the destabilization of the cortical LTP as well as to the blockade of the cooperative maintenance of the weak thalamic LTP (Supplementary Figure 1E). Analysis of the percentage decay of LTP showed that the maintenance of LTP in cortical and thalamic synapses depend on the synthesis of PRPs or its capture through synaptic cooperation (Supplementary Figure 1F).

Since the maintenance of LTP is dependent on this interplay between synaptic tags and the availability of PRPs, to test whether thalamic and cortical synapses compete, we increased the pool of activated synapses, by stimulating a second thalamic input (thalamic W2). Our previous results showed that the time window for thalamic synapses to effectively capture PRPs was around 7.5 minutes (Fonseca, 2013). Thus, we stimulated a second thalamic input (W2), with a weak tetanic stimulation, 7.5 minutes after the first weak thalamic (W1) stimulation. We observed that the LTP in the thalamic W2 was not maintained and that the LTP in the thalamic W1 was destabilized, leading to its decay. Interestingly, the cortical LTP (S3) was not significantly destabilized by this thalamic competition (Figure 1A) and competition was not observed in the absence of W2 stimulation (Figure 1B).

**Figure 1.**
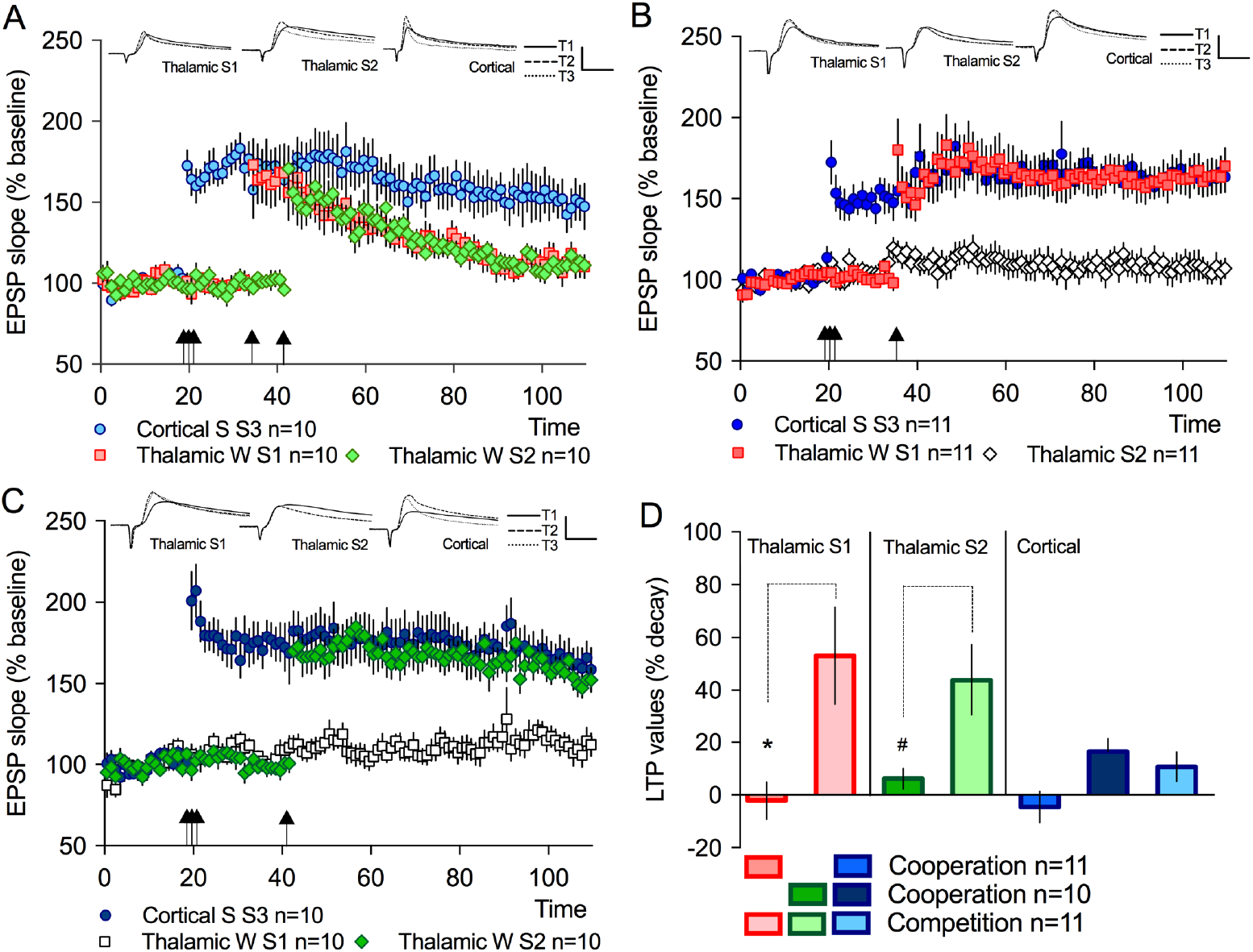
Competition was induced by increasing the pool of activated thalamic synapses. A. Weak stimulation of the Thalamic W2 led to the destabilization of the thalamic W1 LTP, but not of the cortical S3 LTP. B. If the Thalamic W2 was not weakly stimulated, LTP in the cortical S3 and thalamic W1 was maintained, by cooperation. C. Cooperation was also observed if weak thalamic W2 was stimulated 22.5 minutes after cortical S3 strong stimulation. D. The decay of LTP in the cortical S3 was similar in all conditions tested, whereas the Thalamic LTP, in W1 and W2, decayed significantly more if both thalamic inputs were stimulated (*p=0.007; # p=0.01). Inserts represent EPSPs traces before (T1), after LTP induction (T2) and at the end of the recording (T3). Bars: 20 ms and 15mV. Error bars represent SEM, n=number of slices.

To rule out that the decay observed in the thalamic W2, under competition, was due to the time of weak W2 stimulation, we did a cooperation experiment in which the weak stimulation of W1 is not present. Thalamic W2 LTP was still maintained even if stimulated 22.5 minutes after cortical strong stimulation (Figure 1C), showing that at this later time point W2 is still able to capture PRPs. No significant change in LTP decay was observed for the cortical input in all conditions (Figure 1D). These observations suggest that competition results from an imbalance between the availability of PRPs and the pool of activated synapses in a source-to-sink distribution.

### Modulation of the endocannabinoid system alters synaptic competition

We have previously found that inhibiting CB1R extends the time-window of thalamic cooperation (Fonseca, 2013). To test whether modulation of the endocannabinoid system also plays a role in synaptic competition, we did a competition experiment while decreasing or increasing endocannabinoid signalling. We found that pharmacological inhibition of CB1R led to an increase in competition, resulting in the destabilization of LTP in all inputs, thalamic and cortical (Figure 2A; Supplementary Figure S2). Conversely, inhibition of anandamine degradation, by URB597 a specific inhibitor of FAAH, reduced competition and allowed the maintenance of LTP in all stimulated inputs (Figure 2B). Importantly, the inhibition of CB1R by AM281 did not block LTP maintenance if competition was not induced (Figure 2C). Analysis of the percentage decay of LTP showed that URB597 treatment significantly decreased the decay of LTP in both thalamic W1 and W2 whereas AM281 application significantly increased cortical S3 decay under competition (Figure 2D). These results show that PRPs synthesised upon strong cortical stimulation are able to promote the maintenance of all activated synapses if CB1R activation is increased. Since the availability of PRPs is similar in all conditions tested, this observation suggests that CB1R activation modulates the strength of the tag gating the balance towards cooperation, if CB1R activation is increased, or towards competition, if CB1R activation is decreased.

**Figure 2.**
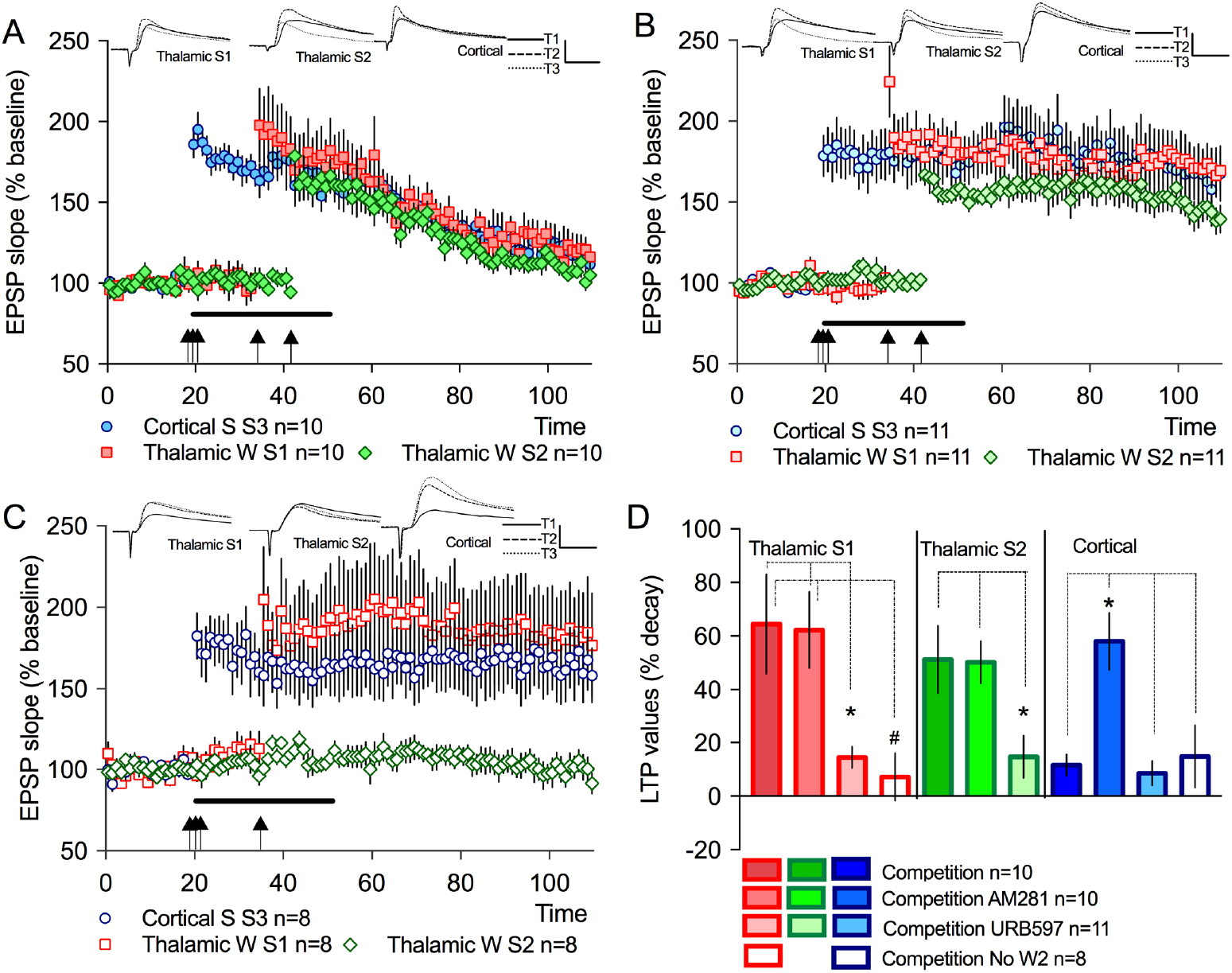
Competition was modulated by the activation of CB1 R. A. Inhibition of CB1R, by AM281application (0.5μM – solid line) led to an increase in competition and to the destabilization of the cortical S3 LTP. B. Conversely, the inhibition of anandamine degradation, by URB597 application (1μM – solid line), led to the maintenance of LTP in all inputs. C. Inhibition of CB1R, by AM281 application (solid line) did not induce competition if the thalamic W2 input was not stimulated. D. The decay of LTP in the strong cortical was significantly higher in the presence of AM281, whereas the thalamic LTP decayed significantly less in the presence of URB597 or when the thalamic W2 was not stimulated. Inserts represent EPSPs traces before (T1), after LTP induction (T2) and at the end of the recording (T3). Bars: 20 ms and 15mV. Error bars represent SEM, n=number of slices.

### Competition depends on the relative time of synaptic stimulation

Since the tag in restricted in time, the sequence of synaptic activation between the multiple inputs will determine whether PRPs are still available, but also if synapses are still capturing PRPs. In our experimental conditions, competition is induced by the stimulation of the second thalamic input that creates an additional pool of activated synapses that capture PRPs. To test whether synaptic competition depends on the time interval of the different input stimulation, we delayed the weak thalamic W2 stimulation to 65 minutes. Based on our previous findings, by increasing the interval between the first and second thalamic stimulation to 30 minutes, we predicted that W1 is no longer destabilized by W2 stimulation. Indeed, we found that synaptic competition is abolished and no destabilization was observed in the thalamic W1 LTP maintenance (Figure 3A).

**Figure 3.**
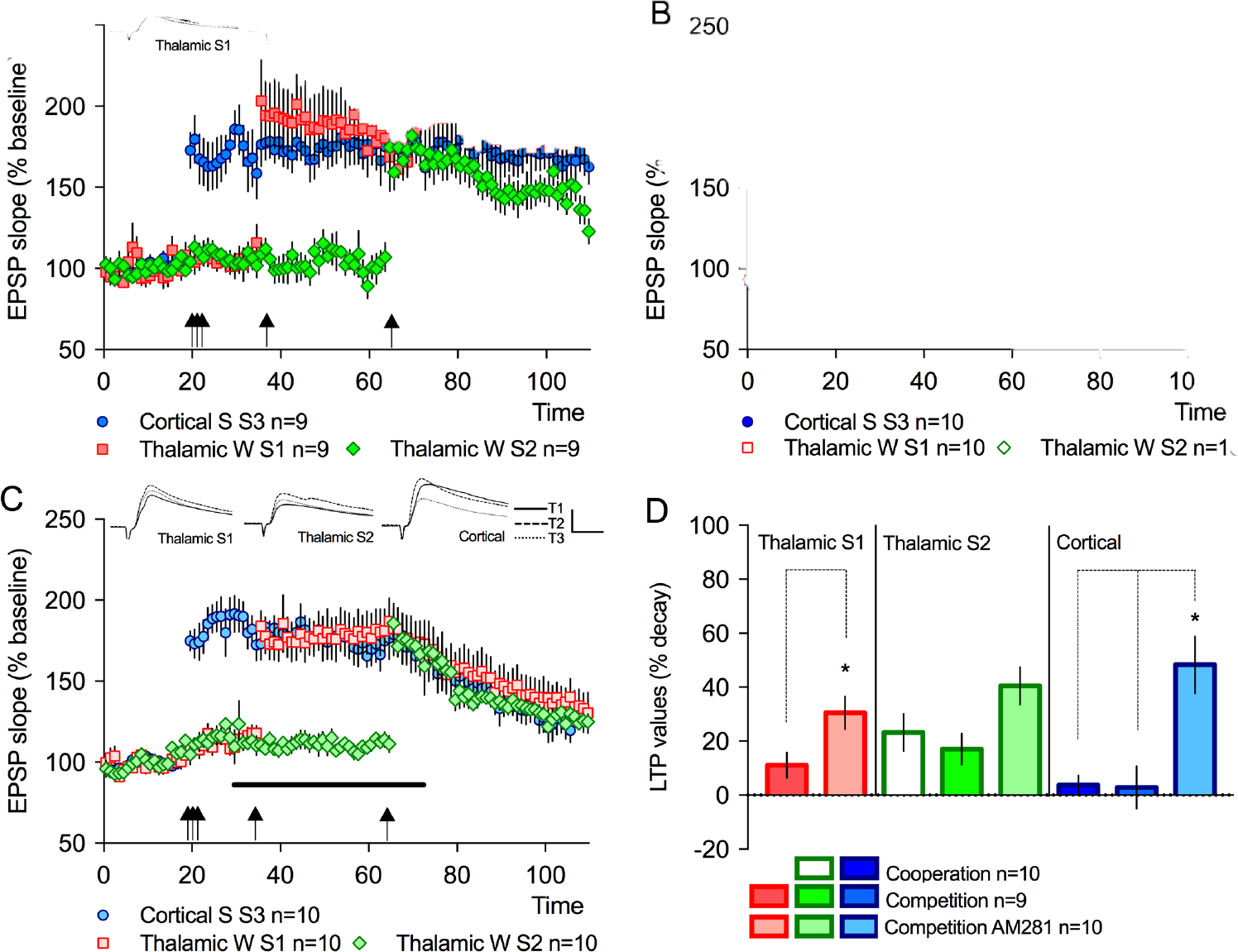
Delaying the stimulation of the thalamic W2 prevents competition. A. If thalamic W2 weak stimulation was delayed to 65 minutes (30 minutes after W1 stimulation) competition was not observed. B. Thalamic W2 was not maintained by cooperation if weak stimulation was delayed to 65 minutes (45 minutes after cortical stimulation). C. Application of AM281 (solid line) increases competition even if thalamic W2 weak stimulation is delayed to 65 minutes, leading to the destabilization of the cortical and thalamic W1 LTP. D. The decay of LTP in the strong cortical and the thalamic W1 was significantly higher in the presence of AM281, whereas the decay of the thalamic W2 did not differ in all conditions. Inserts represent EPSPs traces before (T1), after LTP induction (T2) and at the end of the recording (T3). Bars: 20 ms and 15mV. Error bars represent SEM, n=number of slices.

Interestingly, thalamic W2 LTP was not stabilized, suggesting that PRPs, at this time point, are not sufficient to maintain the thalamic W2 LTP. This interpretation is corroborated by the observation that cooperation also did not occur if strong cortical S3 and weak W2 thalamic stimulation were separated by 45 minutes (Figure 3B). As in the previous experiment, inhibiting CB1 R increased competition. In the presence of AM281, stimulation of thalamic W2 promoted the destabilization of thalamic W1 and cortical S3 LTP (Figure 3C). Weak stimulation of thalamic W2, under CB1R inhibition, significantly increased the LTP decay of the thalamic W1 and cortical S3 LTP (Figure 3D).

### Competition depend on the availability of PRPs

Since competition relies on the balance between the source and the sink of PRPs, then increasing the availability of PRPs, the source, should abolish competition. In our experimental condition, PRPs synthesis was induced by the strong cortical stimulation. To test this, we shortened the time interval between weak thalamic and strong cortical synaptic stimulation. We found that, when thalamic S1 and S2 were stimulated in the close temporal vicinity to cortical strong stimulation, LTP in all activated inputs was maintained (Figure 4A). Once more, CB1R inhibition led to an increase in competition and to the destabilization of LTP in all stimulated inputs (Figure 4B). Reducing the time interval of weak thalamic stimulation in relation to strong cortical LTP, decreased competition, reducing LTP decay. Inhibiting CB1R activation restored competition, suggesting that blocking CB1R created a new imbalance between source and sink (Figure 4C).

**Figure 4.**
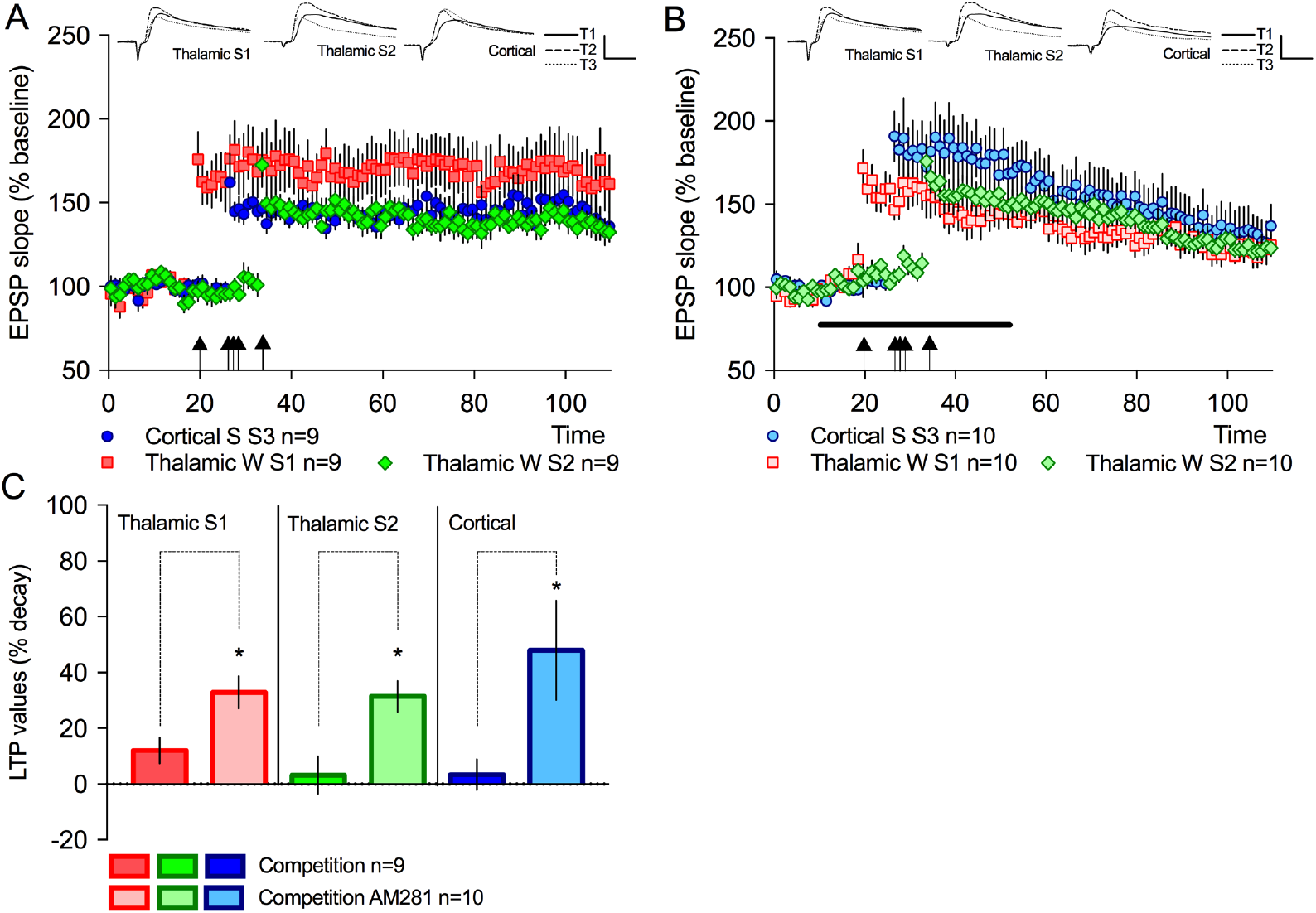
Competition is abolished if PRPs availability is increased. A. Shortening the time interval between the weak thalamic W1 and W2 and the cortical S3 stimulation abolished competition. B. Application of AM281 increased competition and led to the destabilization of LTP in all inputs. C. Percentage decay of LTP for all conditions tested. The decay of LTP is significantly higher for all inputs in the presence of AM281. Inserts represent EPSPs traces before (T1), after LTP induction (T2) and at the end of the recording (T3). Bars: 20 ms and 15mV. Error bars represent SEM, n=number of slices.

Since the inhibition of CB1R increased competition and considering that competition is determined by a source-to-sink balance, then CB1R inhibition may increase the sink (tag) or decrease the source (PRPs). Although CB1R inhibition is associated with a decrease in protein synthesis, by modulation of the mTOR pathway, the observation that inhibiting CB1R promotes cooperation (Fonseca, 2013), does not support the hypothesis that CB1R inhibition decreases the source. An alternative hypothesis is that inhibition of CB1R increases the sink, which goes in line with CB1R inhibition promoting cooperation. Given that CB1R inhibition, in a competitive setting, led to the destabilization of the cortical LTP, we hypothesised that CB1R modulation is higher in thalamic synapses than in cortical synapses. If indeed CB1R preferentially modulates the tag strength of thalamic synapses, then inhibiting CB1R during competition would create an imbalance towards thalamic synapses forcing the cortical synapses to lose. To test this hypothesis, we assessed the modulation of thalamic and cortical synapses by CB1R. CB1R activation is necessary for the induction of the depolarizing-induced suppression of excitation (DSE). This short-term plasticity is associated with a decrease in presynaptic glutamate release induced by transient depolarization (Kodirov et al., 2010). The induction of DSE resulted in a significant decrease in EPSC amplitude both in the thalamic and cortical activated synapse (Supplementary Figure 3A). Inhibition of CB1R with 0.5μM AM281 application blocked the induction of DSE in cortical but not in thalamic inputs (Supplementary Figure 3B), whereas inhibition of CB1R with 1μM AM281 application blocked the induction of DSE in both cortical and thalamic inputs (Supplementary Figure 3C). Since a higher concentration of AM281 was necessary to fully block DSE in thalamic synapses, our results support the hypothesis that the expression, or activity, of CB1R is higher in thalamic inputs.

To further confirm these findings, we analysed the effect of a CB1 R agonist in EPSPs evoked by thalamic and cortical input stimulation. Bath application of Win55,212-2 reduced EPSP slope evoked by thalamic and cortical input stimulation (Supplementary Figure 4A). As for the DSE experiment, a stronger effect of the CB1R agonist was observed in thalamic synapses (Supplementary Figure 4A/B). Analysis of pair pulse facilitation (PPF), evoked by stimulation of thalamic or cortical inputs showed a significant decrease in PPF, suggesting a presynaptic effect of CB1R activation (Supplementary Figure 4C). Taken together, our results suggest that CB1 R activation has a major role in modulating thalamic synapses with a clear impact in restricting both cooperation and competition between cortical and thalamic synapses.

### Decreasing inhibition promotes synaptic cooperation

It is well established that CB1R activation can modulate excitation as well as inhibition (Azad et al., 2003), resulting in an overall decrease in amygdala excitability (Pistis et al., 2004). Thus, one possibility is that the inhibition of CB1 R increases competition by increasing excitability. To test this hypothesis, we used Picrotoxin, an inhibitor of GABA A receptors, to increase excitability in the lateral amygdala during competition. Since decreasing inhibition reduces the threshold of LTP induction, we used the experimental design of late W2 thalamic stimulation, described in Figure 3A, to avoid overlap of picrotoxin with LTP induction. Picrotoxin application (25μM) was restricted to the time interval between the W1 and W2 weak thalamic stimulation (from 35 to 60 minutes). Additionally, we confirmed that weak thalamic stimulation, in this experimental condition, did not led to a maintained LTP (Supplementary Figure 5A). We then tested whether weak stimulation of W2 would promote competition, under decreased inhibition. We did not observe competition, rather, we found that, under reduced inhibition, the weak thalamic W2 was now maintained, despite its late induction (Figure 5A). Co-application of rapamycin with picrotoxin abolished the maintenance of the LTP in the thalamic W2, showing that weak thalamic LTP maintenance occurs through cooperative sharing and capture of PRPs (Supplementary Figure 5B). Taken together, decreasing inhibition did not increase competition but rather allowed the maintenance of the weak thalamic W2, in a protein-synthesis dependent manner. This suggests that picrotoxin increases the availability of PRPs which were captured by the weak thalamic S2. Previous studies have found an association between protein synthesis and neuronal activity, in particular action-potential firing (Fonseca et al., 2006; Karpova, 2006). Since our cells fire more action potentials, under picrotoxin application, we repeated the experiment described above but prevented the cells from firing during the time period of picrotoxin application. Cells were voltage-clamped to −75mV, from 40 to 60 minutes. Preventing neuronal firing abolished the maintenance of the weak thalamic S2 LTP (Figure 5B). In this condition the cooperative maintenance of the thalamic W2 LTP, promoted by picrotoxin, was abolished. Further, the number of spikes negatively correlated with the percentage of LTP decay in the weak thalamic W2 pathway (Figure 5C). This argues for an increase PRPs synthesis induced by spiking, favouring the maintenance of the transient weak thalamic LTP through synaptic cooperation. Inhibition of protein synthesis, at this later time point, had no effect in thalamic W1 and cortical LTP, showing that LTP maintenance at this point was no longer dependent on protein synthesis (Figure 5D).

**Figure 5.**
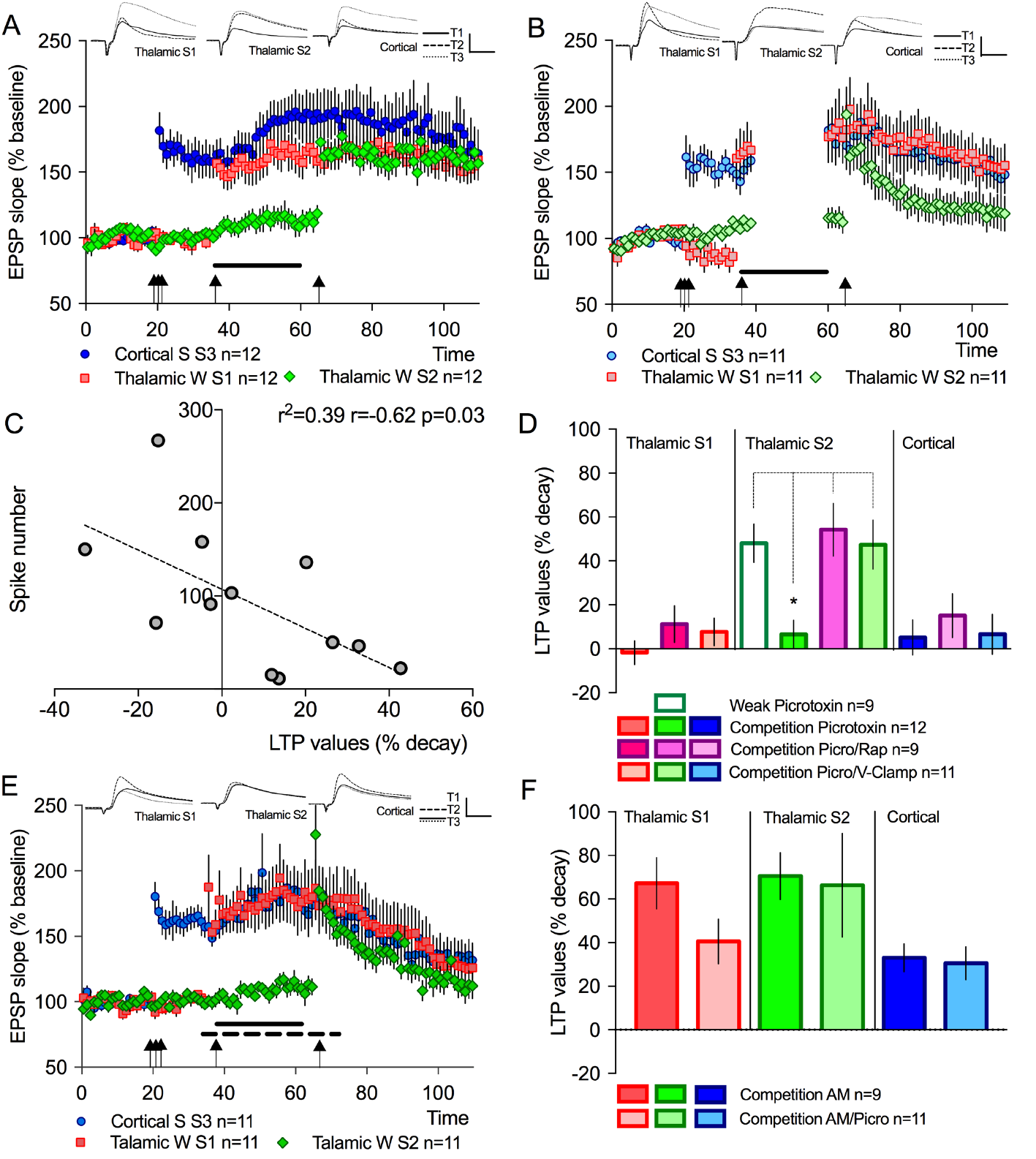
Decreasing inhibition promotes cooperation. A. Decreasing inhibition, by Picrotoxin application (25μM – solid line), allowed the maintenance of LTP in all inputs. B. Maintaining the cells in voltage-clamp mode abolished the cooperative maintenance induced by picrotoxin application. C. Correlation plot of the total number of spikes throughout the recorded time against the percentage decay of the thalamic W2. The significant negative correlation showed that a higher spike number decreases thalamic W2 decay. D. Percentage decay of LTP for all conditions tested. The decay of LTP in the thalamic W2 is significantly lower if slices are treated with picrotoxin, before thalamic W2 stimulation. No significant change in LTP decay was observed in the cortical and thalamic W1. E. Inhibition of CB1R, by AM281 (dashed line), increased competition even if GABA A receptors are also inhibited (picrotoxin – solid line). F. No difference was observed in LTP decay for AM281 or co-application of AM281 with picrotoxin or SCH-50911. Inserts represent EPSPs traces before (T1), after LTP induction (T2) and at the end of the recording (T3). Bars: 20 ms and 15mV. Error bars represent SEM, n=number of slices.

If decreasing inhibition favours an increase in PRPs availability, it is conceivable that it abolishes the increase in competition observed under inhibition of CB1 R. To test this, we did a late competition experiment with the coapplication of AM281 and picrotoxin. The inhibition of CB1R with AM281 covers the induction of LTP in both W1 and W2 thalamic inputs whereas AM281/picrotoxin co-application was restricted to the time window between S1 and S2 thalamic stimulation. In this experimental setting, we still observed competition similar to AM281 application alone (Figure 5E/ Supplementary Figure 5C). A similar result was obtained by co-application of SCH-50911, an inhibitor of GABA B receptors (Supplementary Figure 5D). Analysis showed no differences, in the percentage decay of LTP, between AM281, AM281/picrotoxin and AM281/SCH-50911 co-application (Figure 5F). This shows that the decay in LTP induced by CB1R inhibition, in competition, is not due to an increase in inhibition. Rather, CB1R inhibition creates a new unbalance by increasing thalamic activation.

## DISCUSSION

In this study, we present the first evidence that thalamic and cortical inputs, to the lateral amygdala, interact by synaptic competition. Synaptic cooperation and competition are two forms of heterosynaptic plasticity that occur within large time intervals and result from the synaptic integration of biochemical signals. As described in hippocampal CA3 to CA1 synapses (Fonseca et al., 2004; Sajikumar et al., 2014), competition in the amygdala was observed when the pool of activated synapses was higher than the availability of PRPs. Our results support the hypothesis that synaptic cooperation and competition are determined by a balance between the pool of activated synapses (sink) and the availability of PRPs (source). In our experimental setting, PRPs synthesis was induced by the strong stimulation of the cortical input to the lateral amygdala (sink and source), whereas two thalamic inputs were used as two additional sinks. Because cortical fibers are packed in a bundle we were unable to stimulate two independent cortical pathways, a pre-requisite to assess synaptic cooperation or competition. When a second thalamic input was stimulated with a weak tetanus, sufficient to induce a synaptic tag but not PRPs synthesis, competition led to the decay of a previously induced thalamic LTP (Figure 6A). Interestingly, cortical LTP was not destabilized by thalamic competition. One possibility is that elapsed time consolidated cortical LTP rendering it resistant to disruption by competition. This hypothesis is supported by the observation that inhibiting protein synthesis after cortical strong stimulation had no effect in cortical maintenance. An alternative hypothesis is that the cortical LTP was maintained because the strength of the cortical tag, or the ability of cortical synapses to capture PRPs, was higher than in thalamic synapses. In this scenario, the strength of the tag determines winners and losers. We have previously found, in hippocampal synapses, that the strength of synaptic stimulation correlates with the strength of the tag and with competitive load (Fonseca et al., 2004). Thus, if the thalamic weak stimulation induces a weaker synaptic tag than the strong cortical stimulation, cortical synapses will be able to capture more PRPs, win the competition and maintain LTP.

**Figure 6.**
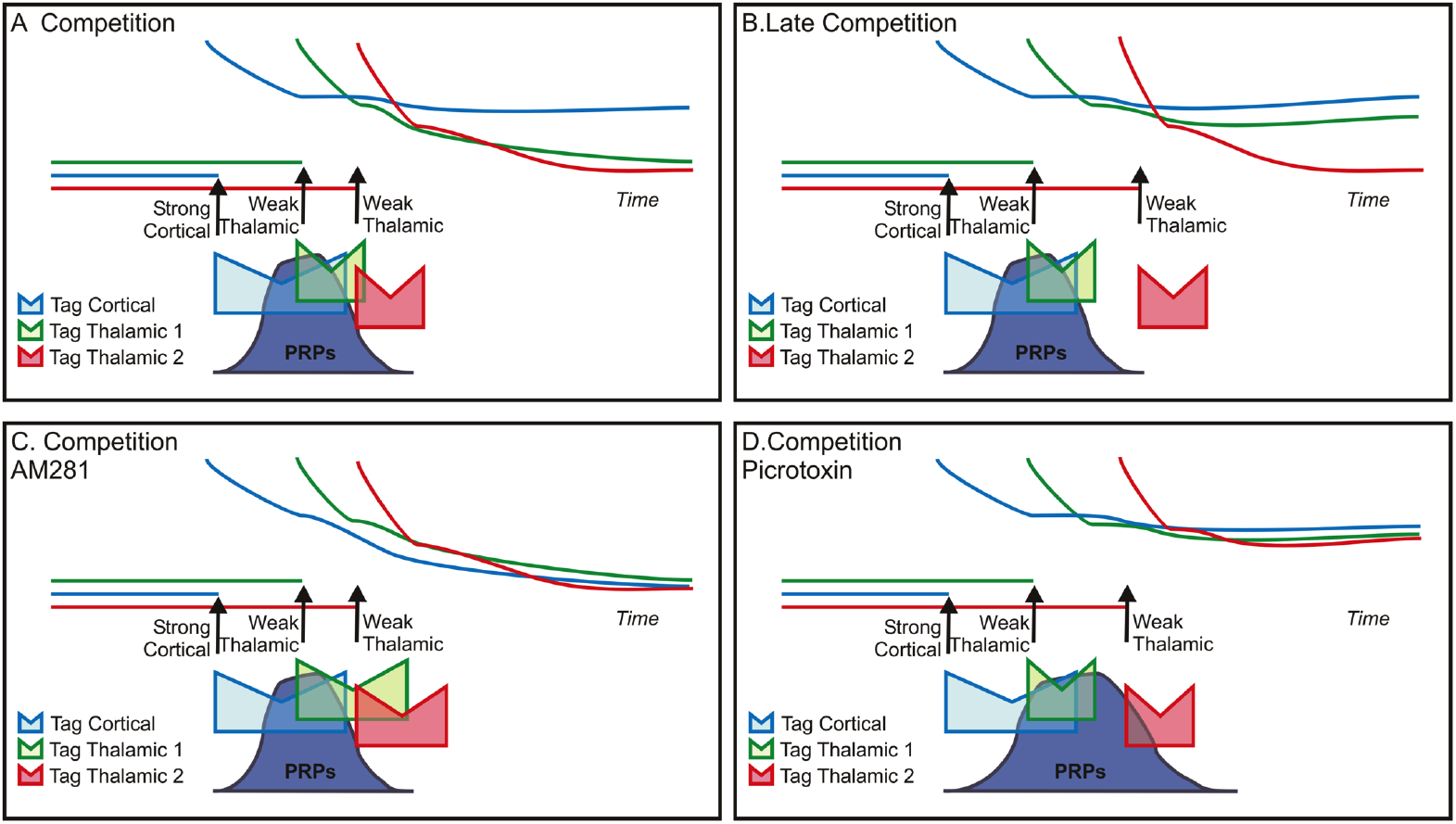
Model of synaptic cooperation and competition. A. In competition, the stimulation of the thalamic W2 leads to the induction of an additional sink that creates an imbalance between the availability of PRPs and the number of tags destabilizing thalamic LTP. B. If the second thalamic stimulation is delayed, the strong cortical and the weak thalamic W1 are stabilized and resistant to competition. C. Inhibition of CB1R increases the duration of the tag promoting competition. D. Decreasing inhibition promotes PRPs synthesis, decreasing the competitive load and promoting cooperation.

The hypothesis that the degree of synaptic activation correlates with the strength of the tag is supported by a previous study showing that suspending synaptic activation abolishes PRPs capture and plasticity maintenance (Szabó et al., 2016). It also goes in line with our previous finding, that inhibiting CB1R extends the time-window in which thalamic synapses can capture PRPs (Fonseca, 2013). If CB1R activation acts as a negative feedback signal, decreasing excitatory synaptic transmission, then inhibiting CB1R increases synaptic activation and therefore the strength of the synaptic tag. In this scenario, inhibiting CB1R, in a competitive setting, would lead to an increase of the tag (sink) and to the competitive load. This is indeed what we observed under CB1 R inhibition, where competition is promoted (Figure 6C). In this functional model, the strength of the tag is modulated by on-going synaptic activation, determining whether a subsequent synaptic stimulation will be maintained by cooperation or destabilized by competition. Our data show that in amygdala synapses, activation of presynaptic CB1R decreases the tag and contribute to a restriction in the time window of cooperation and competition.

It is also possible that inhibition of CB1R decreases the availability of PRPs promoting competition. Indeed, two previous reports have shown that CB1R inhibition downregulates the activity of the mTOR pathway (Busquets-Garcia et al., 2013; Puighermanal et al., 2013). However, since CB1R inhibition promotes cooperation (Fonseca, 2013), it is highly unlikely that it decreases PRPs synthesis. Also, we did not observe any effect on LTP maintenance, under AM281 application, when the thalamic S2 input was not stimulated.

Competition is also modulated by the availability of PRPs, which in turn, is time and activity-dependent. In the case where weak thalamic stimulation was temporally closer to the strong cortical stimulation, competition was not observed, suggesting that the initial availability of PRPs was sufficient to maintain LTP in all activated synapses. Interestingly, the overall neuronal activity can also modulate PRPs synthesis. When inhibition was blocked, PRPs availability increased, allowing the cooperative maintenance of a weak thalamic input at much later time points. The link between neuronal excitability and the synthesis of PRPs has been widely reported (Barco et al., 2002; Ehlers, 2003; Han et al., 2008). Our results suggest that inhibition in the amygdala also modulates the synthesis of PRPs, restricting synaptic cooperation and thus, promoting competition. It is also interesting that reducing inhibition, by either GABA A or B receptor inhibition, did not prevent the increase in competition induced by the CB1R blockade. This finding rules out the possibility that the increase in competition, observed under CB1R inhibition, was a reflection of an increase in network inhibition.

It is now clear that cooperation and competition are mechanisms involved in memory acquisition and maintenance (Cai et al., 2016; Kastellakis et al., 2015; Rashid et al., 2016; Tonegawa et al., 2015). However, the rules by which these two processes are orchestrated are still unclear. Our results show that the time interval between the occurrence of two events will determine whether synaptic cooperation or competition is favoured. Recent studies have also presented compelling evidence that neuronal excitability determines which neurons are recruited to participate in the encoding of a particular memory (Yiu et al., 2014). Our data are consistent with this hypothesis and extends it in a significant manner. By activating CB1R, pyramidal cells in the amygdala regulate their intrinsic excitability reducing the probability that subsequent events are linked. Taken together, our results set up a strong conceptual model, built from a detailed analysis of synaptic plasticity in the amygdala, that provides clear-cut predictions regarding the rules of memory acquisition and maintenance. Given that endocannabinoid signalling is involved in anxiety, our results support the hypothesis that by limiting cooperation and competition, the activation of cannabinoid receptors may restrict fear generalization, one of the mechanisms underlying the development of anxiety disorders.

## FUNDING

This work is supported by a grant from the Portuguese Research Council (Fundação para a Ciência e Tecnologia – 02/SAICT/2017/030772) and a Young Investigator Grant from the Brain and Behaviour Research Foundation (NARSAD grant number 25118). Rosalina Fonseca is supported by an FCT Investigator grant (IF/01359/2014) and Natália Madeira is supported by a PhD fellowship (SFRH/BD/130911/2017).

## ACKNOWLEDGMENTS

We are grateful to Dr. Rita Teodoro for helpful comment on the manuscript. We gratefully thank Dr Tobias Bonhoeffer and Dr Volker Staiger for providing software for the electrophysiology experiments.

