## Supplementary figures for "Temporal gating of synaptic competition in the lateral amygdala by cannabinoid receptor modulation of the thalamic input"

**This PDF file includes:**

Supplementary Figures S1 to S5

Supplementary Figures

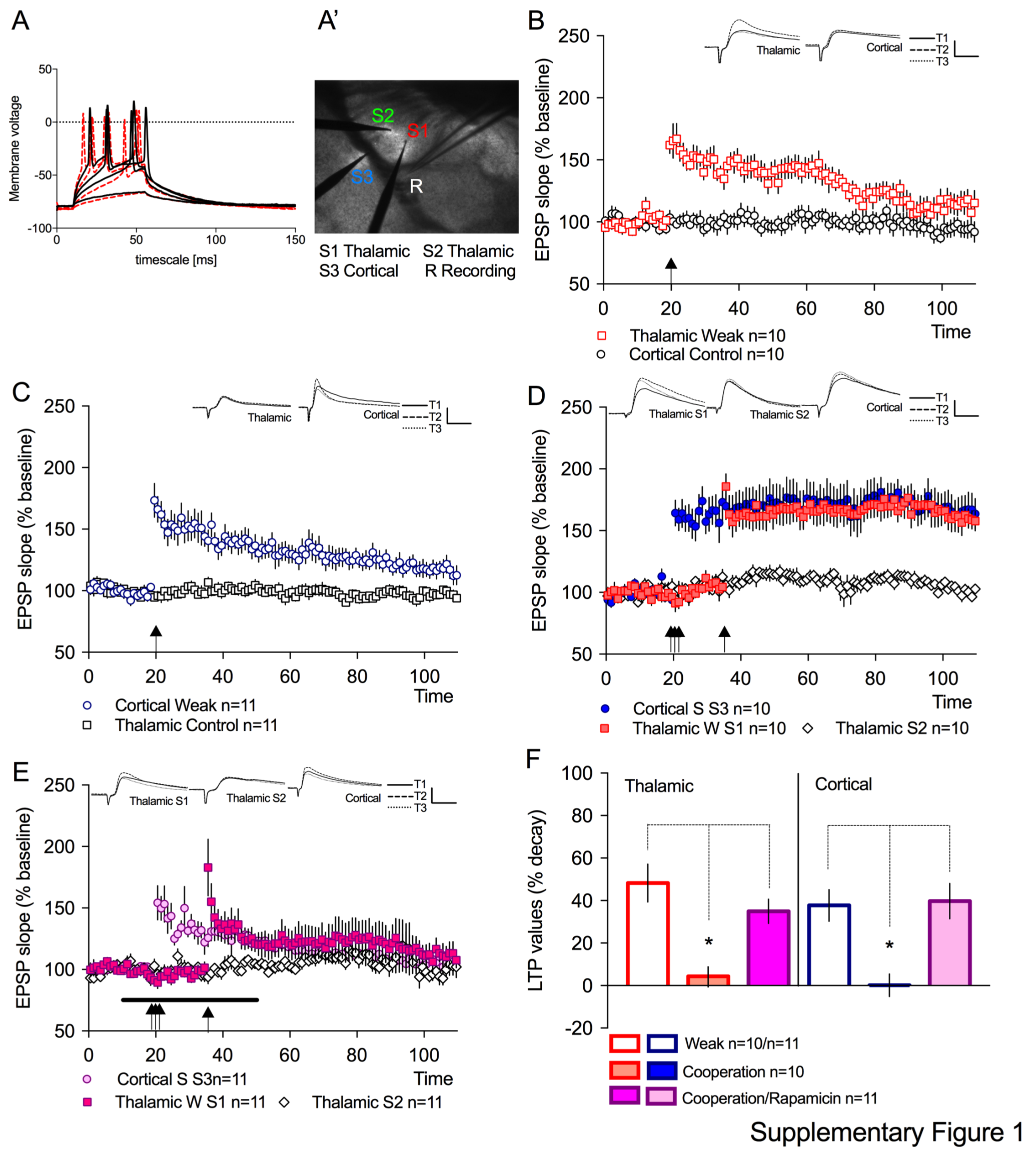


**Figure S1. Cooperation in the amygdala requires *de novo* protein-synthesis.** A. Voltage responses of a LA pyramidal neuron cell in response to steps of depolarizing current injections at the beginning (black) and end of the recording (red). A'. Positioning of the stimulating electrodes in the external capsule (cortical S3), the internal capsule (thalamic W1 and W2) and the recording electrode (R). B. Weak stimulation of the thalamic input induced a transient form of LTP (Thalamic W1). No change was observed in basal synaptic transmission (cortical control). C. Weak tetanic stimulation of the cortical input (cortical weak) also induced a transient LTP. No change was observed in basal synaptic transmission (thalamic control). D. Induction of a maintained LTP, by a strong stimulation of the cortical input (Cortical S3) led to the maintenance of LTP in the thalamic input (Thalamic W1). No change was observed in basal synaptic transmission (thalamic control). E. Application of Rapamycin (1μM) blocked the maintenance of the LTP in the cortical (Cortical S3) and also the maintenance of the weak thalamic LTP (Thalamic W1). Rapamycin was applied for 40 minutes starting 10 minutes prior to strong cortical stimulation. Blocking protein synthesis did not interfere with basal synaptic transmission (thalamic control). F. Percentage of LTP decay for all conditions tested. The decay of LTP in the strong cortical was significantly higher in the presence of rapamycin or when cortical synapses were weakly stimulated (* p<0.01). Thalamic LTP decayed significantly less when strong cortical stimulation preceded weak thalamic stimulation (* p<0.015). Inserts represent EPSPs traces before (T1), after LTP induction (T2) and at the end of the recording (T3). Bars: 20 ms and 15mV. Error bars represent SEM, n=number of slices.

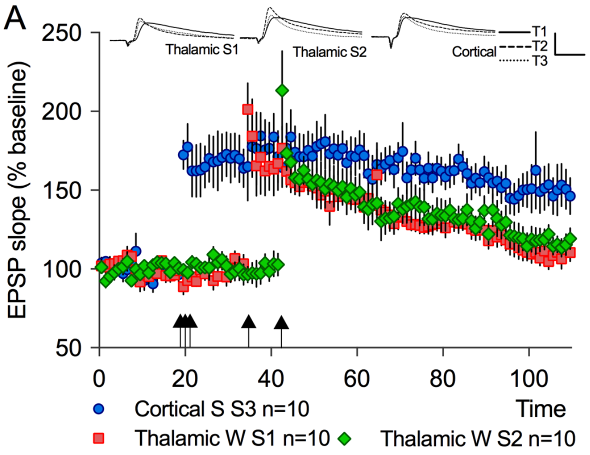


Supplementary Figure 2

**Figure S2. Competition.** A. Similar experiment as depicted in Figure 1A. Data used for group comparison in Figure 2D. Weak stimulation of the thalamic W2 input induced competition leading to the decay of the thalamic W1but not of the cortical

S3. Inserts represent EPSPs traces before (T1), after LTP induction (T2) and at the end of the recording (T3). Error bars represent SEM, n=number of slices.

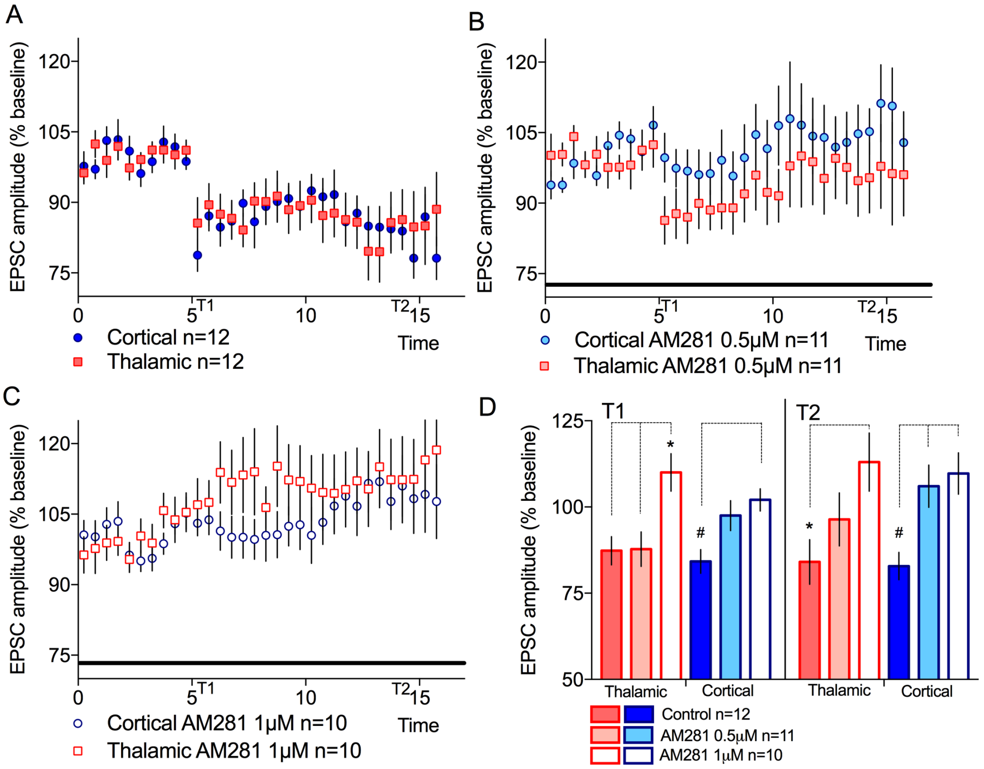


Supplementary Figure 3

**Figure S3. Inhibition of Depolarizing-suppression of excitation (DSE) in the thalamic input, required higher concentration of AM281.** A. Voltage-clamp recordings of EPSCs in amygdala pyramidal cells, evoked by stimulation of the thalamic and cortical inputs. Depolarizing cells to 0mV for 10 seconds, led to a sustained decrease in EPSC amplitude in cortical and thalamic inputs B. Inhibition of CB1 receptors with 0.5μM AM281, blocked DSE induction in the cortical input, but did not fully blocked DSE at the thalamic input. C. Increasing AM281 concentration to 1μM, fully blocked DSE in both cortical and thalamic inputs. D. Average EPSC amplitude for the two time-windows analyzed: T1=5-7 minutes (*p<0.01 #p=0.01) and T2=13-15minutes (*p<0.04 # p<0.01). Error bars represent SEM, n=number of slices.

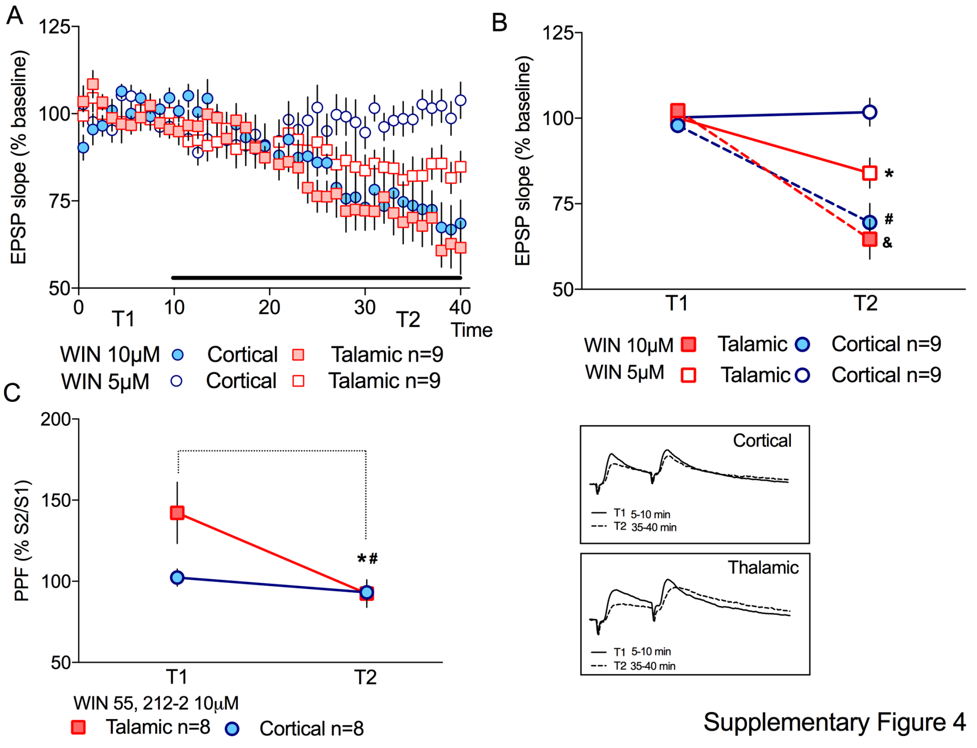


Supplementary Figure 4

**Figure S4. Application of an agonist of CB1 receptors decreased excitatory transmission in thalamic and cortical inputs to the amygdala.** A. Application of Win55,212-2, a CB1R agonist, led to a decrease in EPSP slope, both in thalamic and cortical evoked synaptic responses. Win55,212-2 was applied for 30 minutes starting 10 minutes after the starting of the recording. B. Analysis of the decrease in EPSP slope. Cortical evoked responses were not affected by 5µM but significantly decreased by the application of 10µM of the agonist (Friedman ANOVA *p=0.03; # p=0.004; & p=0.03; T1=5-10min T2=35-40). C. Activation of CB1R by 10µM of Win55,212-2 led to a decrease in pair-pulse facilitation (PPF) in thalamic evoked EPSP (Friedman ANOVA Thalamic *p=0.03; Cortical # p=0.04). Insert represent EPSPs evoked by paired stimulation with an interval of 30ms, before (T1) and after CB1R agonist (T2). Error bars represent SEM, n=number of slices.

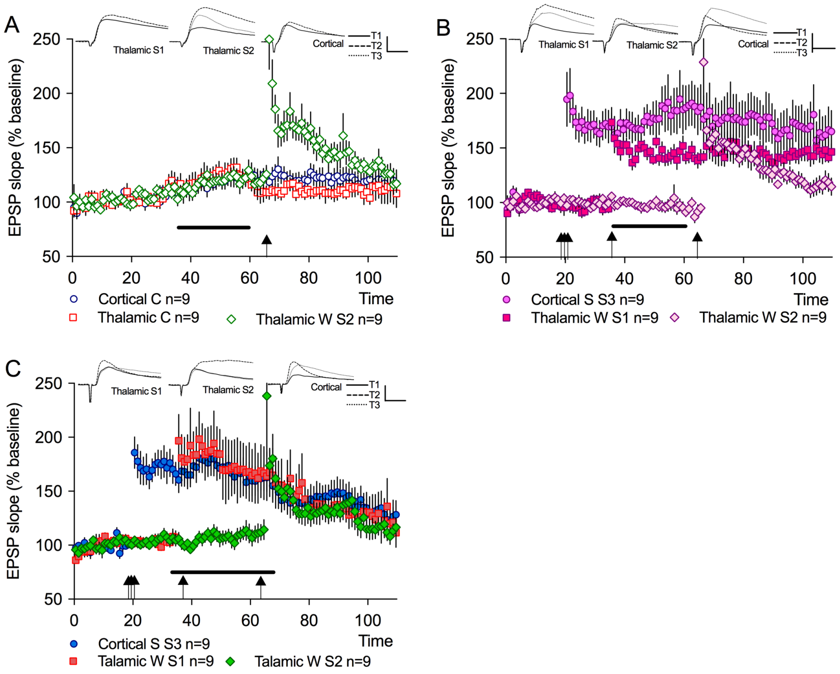


Supplementary Figure 5

**Figure S5. Effect of inhibition in synaptic competition.** A. Weak stimulation of the thalamic W2 input led to a transient LTP, even if slices were treated with Picrotoxin (25μM) for 25 minutes prior to LTP induction (from 35 to 60 minutes – solid line). B. Co-application of Rapamycin with Picrotoxin (solid line) blocked the cooperative maintenance of the weak stimulated thalamic W2. Rapamycin application did not prevent the maintenance of the LTP in the thalamic W1 nor in the cortical S3. C. Inhibition of CB1R, by AM281 application, increased competition. Similar experiment as depicted in Figure 5A. Data used for group comparison in Figure 5F. D. Inhibition of GABA B receptors by the application of SCH-50911 (10μM) did not prevent the increase in competition induced by concomitant inhibition of CB1R (solid line represent AM281 application, dashed line SCH-50911). Data used for group comparison in Figure 5F. Insert represent EPSPs traces before (T1), after LTP induction (T2) and at the end of the recording (T3). Error bars represent SEM, n=number of slices.
